## Supplemental Note for "High-resolution detection of copy number alterations in single cells with HiScanner"

### Supplemental Information: High-Resolution Detection of Single Cell Copy Number Alterations Using HiScanner

#### 1 Supplemental Note: BAF refinement and the statistical test for LOH

Several factors affect read coverage in a genomic region: sequence-related biases, notably GC content and mappability; amplification noise such as allelic imbalance and dropout; and the inherent copy number state.

First reported by Dohm *et al* [1], GC content bias describes the dependency between the proportion of guanine and cytosine bases in a region and the fragment count. This bias persists for bins as large as 100 kb even in bulk sequencing data [2]. While the overall pattern of GC dependency remains consistent, the magnitude of the effect may differ considerably across samples, even those that are matched [3]. CNA callers typically correct this bias by modeling the GC content-read depth relationship by locally estimated scatterplot smoothing (loess) or by using matched normal. Notably, GC bias is expected to exert similar effects on a single position for both homologs because nucleotide content is—in the absence of many mutations—generally similar between the homologs. Although HiScanner also corrects for GC bias using regression, it first leverages the uncorrected read counts to perform the LOH test described below.

Another challenge for single-cell CNA analysis is handling imbalanced amplification between alleles, a result of independent amplification of homologs [4]. The most extreme imbalance, allelic dropout artifacts, is difficult to distinguish from a biological copy number loss. Previous single-cell CNA callers reduce the rate at which amplification artifacts are misidentified as CNAs by averaging them out using large bin sizes and post-hoc filtering of small CNAs; but this approach reduces or eliminates sensitivity for small CNAs. In addition to false positive CNAs, dropout artifacts can also impact BAF estimation by producing bins with spurious BAF=0 or BAF=1 measurements.

HiScanner introduces an efficient BAF estimation procedure that reduces the effect of dropout artifacts by exploiting read depth fluctuations caused by GC bias. In the first step, we differentiate between the case of LOH, for which BAF=0 or 1 is correct, and allelic dropout artifacts, for which BAF=0 or 1 will be observed incorrectly. The property underpinning this test is that the effect of local GC bias (which affects both alleles) in regions without LOH tends to be greater than allelic imbalance between alleles. As a result, changes in total read depth are split in near equal proportions between the two alleles most of the time, evidenced by a BAF distribution centered at 0.5 despite large variances in total read depth (Supplementary Figure 2a). In regions with LOH—where only one allele is present—BAF oscillates between 0 and 1 due to imperfect phasing of gHETs (Supplementary Figure 2d). Importantly, we assume that true LOH events typically span regions larger than most allelic dropout event (i.e., the optimal bin size inferred from the data). We hypothesized that a shift operation, in which shifted BAF' is computed using allele-specific counts from nearby bins (rather than the same bin), would create a different distribution than standard BAF when there is no LOH, but have little to no effect when LOH is present (Supplementary Figure 2b,e). The difference between distributions of standard and shifted BAFs is detected by a Kolmogorov-Smirnov test and, if present, indicates LOH (Supplementary Figure 2c,f).

In more detail, let ai and bi be the haplotype A- and haplotype B-specific read depths at the ith bin in a CNA candidate segment and $BAF_i = \frac{b_i}{a_i+b_i}$. The shifted BAF, which we denote BAF', uses the allele-specific read depth for haplotype A from a bin shifted by k bins: $a_{i+k}; k = 2$ is the default. The shifted BAF is therefore $BAF_i' = \frac{b_i}{a_{i+k}+b_i}$. Both standard BAF and shifted BAF' are mirrored around 0.5 (i.e., $BAF = min(BAF, 1 - BAF)$). When there is no LOH, the difference between ai and ai+k is relatively large due to GC-bias fluctuation, and the shifted BAF is pushed toward 0 (Supplementary Figure 2a vs. Figure 2b). However, when LOH occurs, $a_i = a_{i+1}$ (unless there is a phase-switch error), and the shifted BAF distribution resembles the standard BAF distribution (Supplementary Figure 2d vs. Figure 2e). The effect of phase-switch errors on the shifted BAF distribution is minor since $a_i \neq a_{i+1}$ only for the bin where the error occurred.

For each segment, HiScanner builds the BAF and BAF' distributions and then detects LOH by performing a Kolmogorov-Smirnov (KS) test: if the distributions do not significantly differ, LOH is detected (Supplementary Figure 2f), and the segment is assigned a BAF=0. If the distributions do significantly differ, indicating non-LOH (Supplementary Figure 2c), HiScanner proceeds to the second step: peak calling. Peak calling is performed by applying `scipy.signal.find_peaks` to a histogram of the BAF distribution. The BAF corresponding to the largest non-zero peak–to mitigate the influence of dropout artifact bins with BAF=0 on the segment-wide BAF estimate–is the final segment-wide BAF.

#### 2 Supplementary Data: Simulated ground truth CNA coordinates

The simulated ground truth CNA coordinate are provided in the Source Data. Files are named according to the convention [cn_state].[size].bed, where [cn_state] indicates the event type, and [size] indicates the size of the simulated event in base pairs. The simulations cover a geometric progression of sizes ranging from 200Kb to 5Mb, providing a comprehensive dataset for evaluating the sensitivity and precision of CNV detection methods across different scales of genomic alterations. Each BED file contains the genomic coordinates of simulated CNAs at the specified size, allowing for controlled performance assessment against known ground truth events.

### 3 Supplementary Figures and Tables

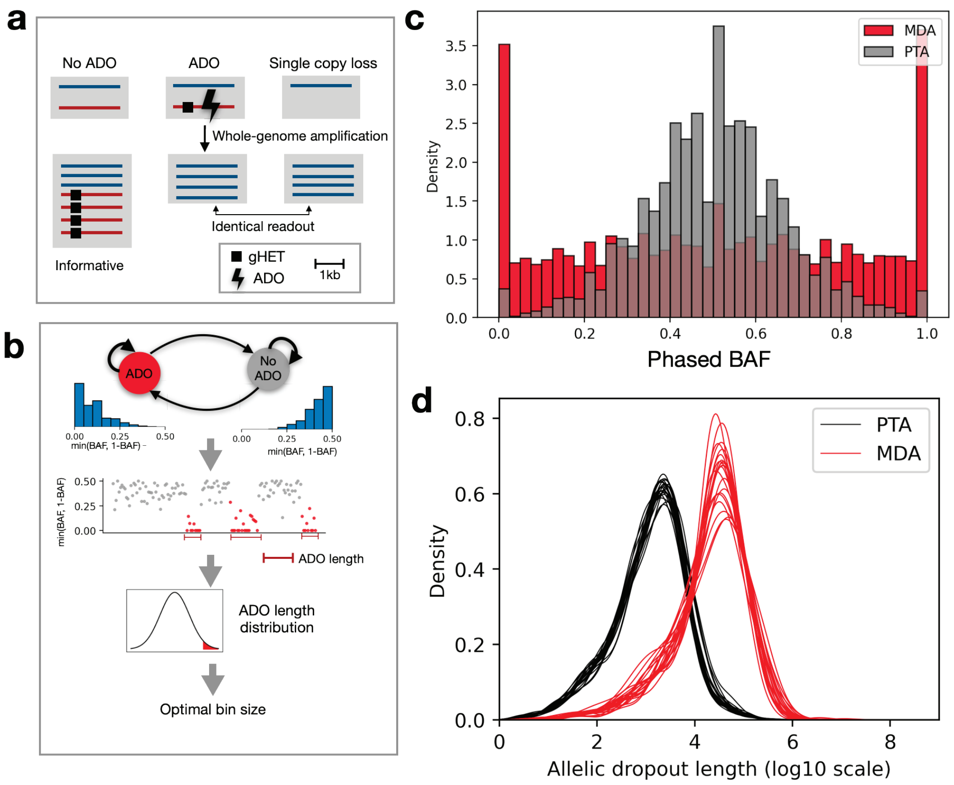


**Supplementary Figure 1** Allelic dropout analysis. **a.** Patterns of scWGA readout in three scenarios: non-ADO, ADO, and single copy loss, in a diploid genome. gHET: germline heterozygous SNP. **b.** The HMM consists of two hidden states: non-ADO and ADO, with distinct emission probability distributions, with mirrored BAF centered at 0.5 and 0, respectively. HMM's prediction of ADO states at individual gHET sites are then used to find length of ADO events and choose the bin size. **c.** BAF distribution of two oligodendrocytes that underwent either PTA or multiple displacement amplification (MDA). **d.** ADO length distribution of the PTA oligodendrocyte (black) and the MDA oligodendrocyte (red). Lines correspond to ADO density in individual autosomes.

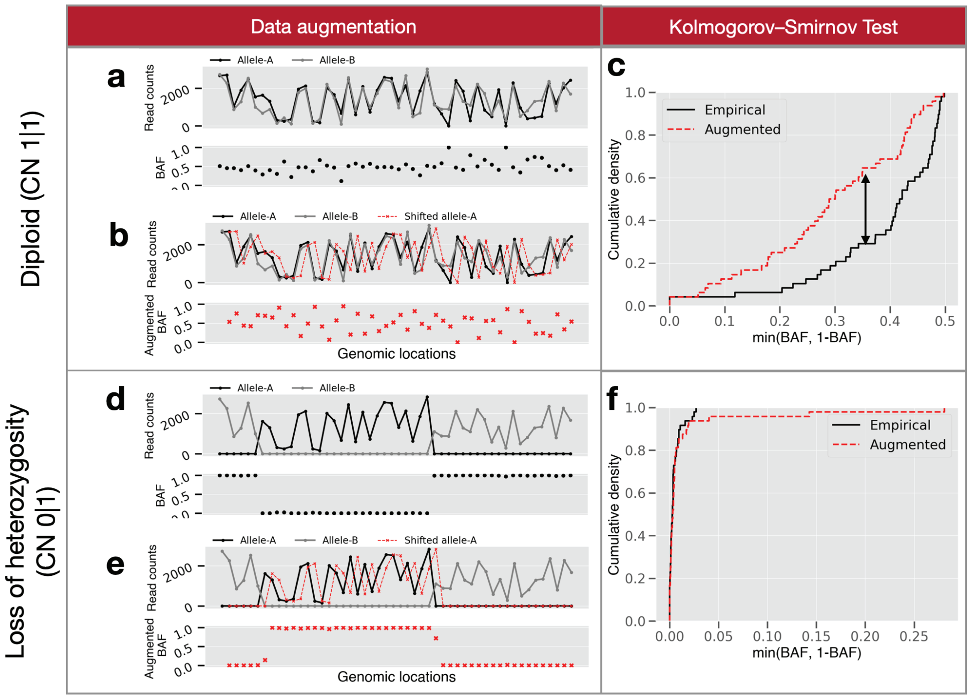


**Supplementary Figure 2** Detailed illustration of HiScanner's LOH test. During the data augmentation phase, read counts corresponding to allele A are shifted rightward by 1 step (two bins in this illustration). **a.** Original read counts for alleles A and B, with corresponding BAF values shown below. **b.** Demonstration of the shift operation with allele A counts shifted rightward, allowing for the calculation of an augmented BAF (shown in red). **c.** The K-S test comparing empirical and augmented BAF distributions for the diploid heterozygous case shows significant difference between the distributions. In this condition, there is noticeable BAF variation from the expected 0.5 value, attributable to differential GC content influence across genomic regions. **d.** Original read counts in a region with loss of heterozygosity, showing the characteristic pattern of predominantly one allele. **e.** The shift operation applied to the LOH region. **f.** The K-S test for the LOH scenario shows minimal difference between empirical and augmented BAF distributions. In this case, the BAF exhibits a mirrored pattern but retains similar absolute deviations from 0.5, barring locations affected by phase switch errors.

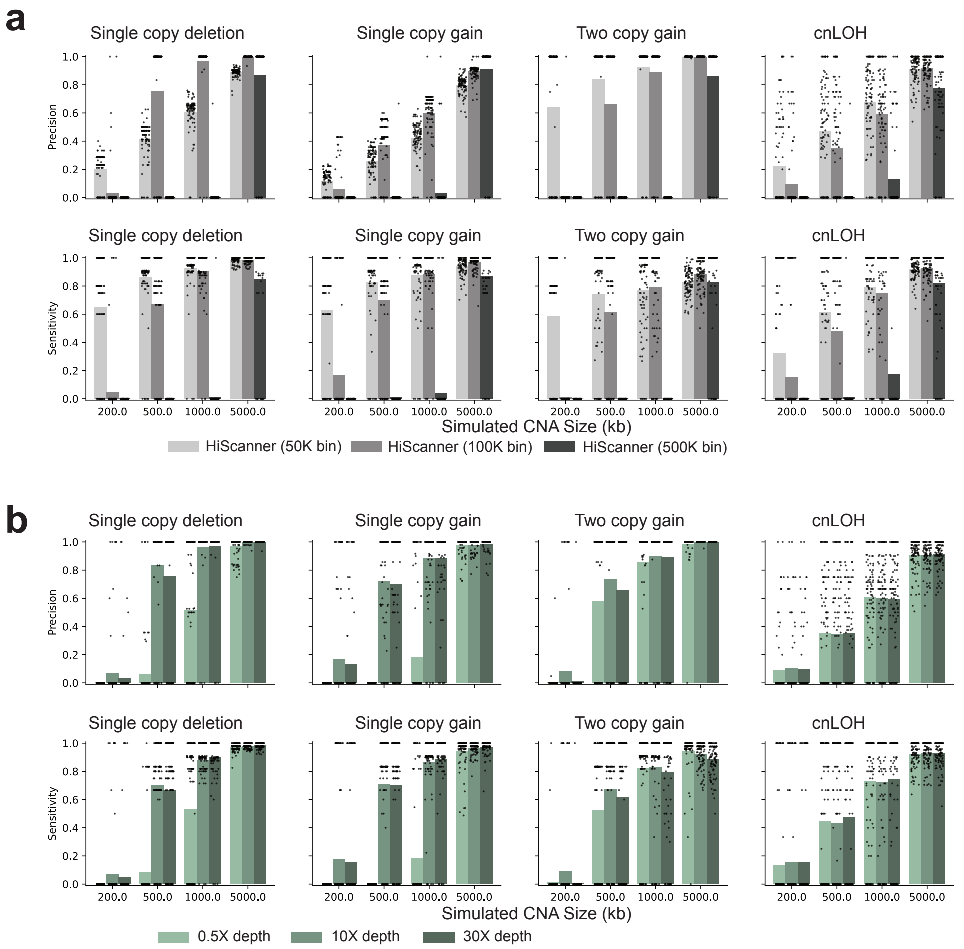


**Supplementary Figure 3** HiScanner performance across various simulated read depths and bin sizes. **a.** Impact of bin size (50Kb, 100Kb, and 500Kb) on precision and sensitivity for detecting different CNV types at 30X coverage. **b.** Effect of sequencing depth (0.5X, 10X, and 30X) on detection performance using 100Kb bins. For each subplot, the x-axis shows CNV size in kilobases and the y-axis shows either precision (top) or sensitivity (bottom). CNV types tested include single copy deletions, single copy gains, two copy gains, and copy-neutral loss of heterozygosity (cnLOH).

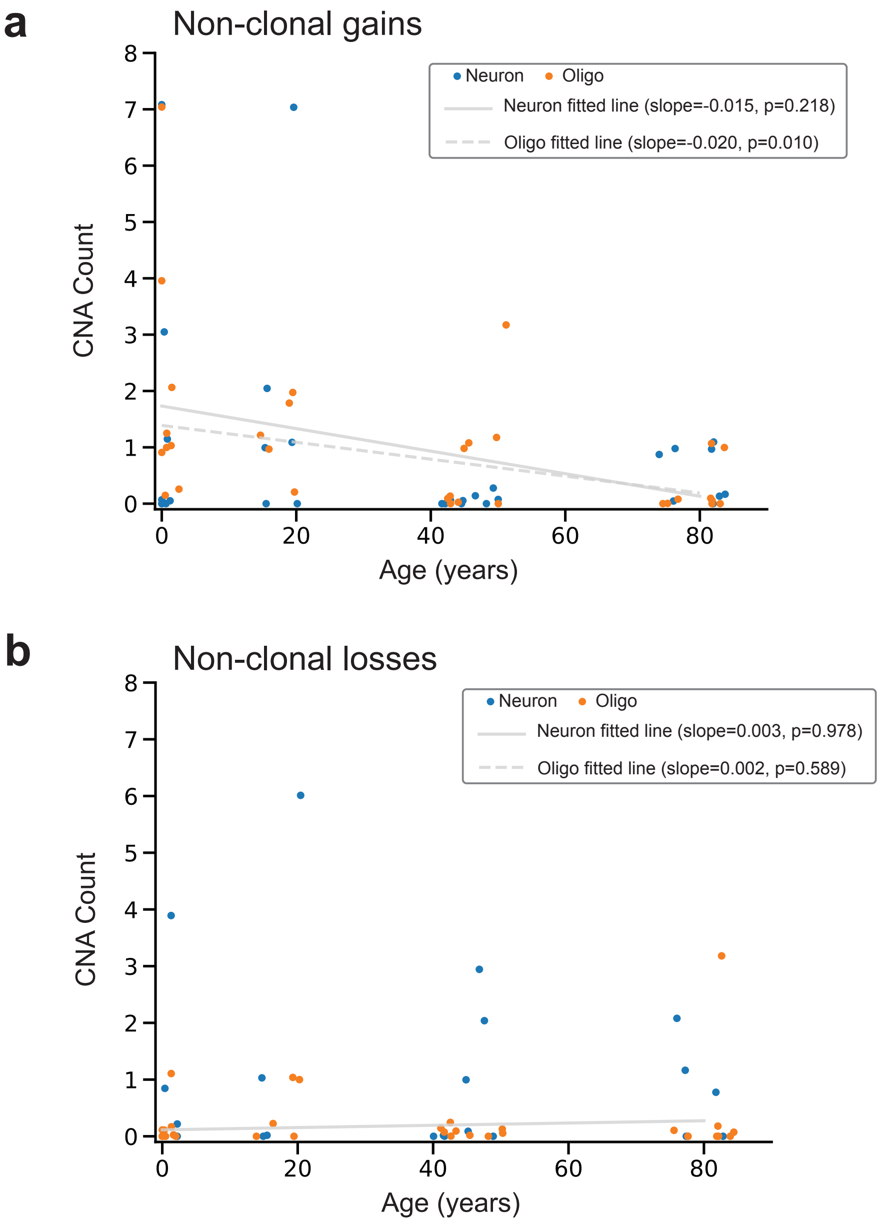


**Supplementary Figure 4** Neuronal and oligodendrocyte CNV burden across age. **a.** Non-clonal losses per cell across different age groups and cell types. **b.** Non-clonal gains per cell across different age groups and cell types. In oligodendrocytes, a weak (slope = -0.02), marginally significant (Bonferroni-adjusted p-value 0.04) negative correlation for private gains was observed, but other tests did not attain statistical significance.

**Supplementary Table 1** False discovery rate (FDR) of the high-depth PTA human brain call set using different size thresholds.

| Size filter | Single copy loss FDR | Single copy gain FDR | Two copy gain FDR | cnLOH FDR |
|---|---|---|---|---|
| No filter | 21.46% | 21.63% | 25.32% | 46.32% |
| 0.25Mb | 15.21% | 16.18% | 19.18% | 42.65% |
| 0.3Mb | 15.21% | 16.18% | 19.18% | 42.65% |
| 0.4Mb | 10.01% | 13.65% | 14.92% | 39.36% |
| 0.5Mb | 7.68% | 12.09% | 12.71% | 36.07% |
| 1.0Mb | 1.17% | 4.10% | 3.00% | 18.49% |
| 2.0Mb | 0.67% | 2.30% | 1.00% | 12.90% |
| 3.0Mb | 0.15% | 1.59% | 0.33% | 11.12% |
| 4.0Mb | 0.15% | 1.43% | 0.00% | 9.14% |
| 5.0Mb | 0.07% | 1.43% | 0.00% | 8.42% |

**Supplementary Table 2** PTA brain cells analyzed in this study. All of the cells were amplified with PTA.

| Cell ID | Sample ID | Cell Type | Age (year) |
|---|---|---|---|
| 936-Oligo-4 | 936 | oligodendrocyte | 49.2 |
| 936-Oligo-5 | 936 | oligodendrocyte | 49.2 |
| 936-Oligo-6 | 936 | oligodendrocyte | 49.2 |
| 936PFC-A | 936 | neuron | 49.2 |
| 936PFC-B | 936 | neuron | 49.2 |
| 936PFC-C | 936 | neuron | 49.2 |
| 1278-Oligo-5 | 1278 | oligodendrocyte | 0.4 |
| 1278-Oligo-7 | 1278 | oligodendrocyte | 0.4 |
| 1278-Oligo-8 | 1278 | oligodendrocyte | 0.4 |
| 1278BA9-A | 1278 | neuron | 0.4 |
| 1278BA9-B | 1278 | neuron | 0.4 |
| 1278BA9-C | 1278 | neuron | 0.4 |
| 4638-Oligo-4 | 4638 | oligodendrocyte | 15 |
| 4638-Oligo-5 | 4638 | oligodendrocyte | 15 |
| 4638-Neuron-4 | 4638 | neuron | 15 |
| 4638-Neuron-5 | 4638 | neuron | 15 |
| 4638-Neuron-6 | 4638 | neuron | 15 |
| 4643-Oligo-5 | 4643 | oligodendrocyte | 42 |
| 4643-Oligo-6 | 4643 | oligodendrocyte | 42 |
| 4643-Oligo-8 | 4643 | oligodendrocyte | 42 |
| 4643-Neuron-3 | 4643 | neuron | 42 |
| 4643-Neuron-4 | 4643 | neuron | 42 |
| 4643-Neuron-6 | 4643 | neuron | 42 |
| 5087-Oligo-4 | 5087 | oligodendrocyte | 44.9 |
| 5087-Oligo-5 | 5087 | oligodendrocyte | 44.9 |
| 5087-Oligo-7 | 5087 | oligodendrocyte | 44.9 |
| 5087PFC-A | 5087 | neuron | 44.9 |
| 5087PFC-B | 5087 | neuron | 44.9 |
| 5087PFC-C | 5087 | neuron | 44.9 |
| 5219-Oligo-5 | 5219 | oligodendrocyte | 76 |
| 5219-Oligo-6 | 5219 | oligodendrocyte | 76 |
| 5219-Oligo-7 | 5219 | oligodendrocyte | 76 |
| 5219-Neuron-2 | 5219 | neuron | 76 |
| 5219-Neuron-4 | 5219 | neuron | 76 |
| 5219-Neuron-5 | 5219 | neuron | 76 |
| 5559-Oligo-3 | 5559 | oligodendrocyte | 19.8 |
| 5559-Oligo-5 | 5559 | oligodendrocyte | 19.8 |
| 5559-Oligo-8 | 5559 | oligodendrocyte | 19.8 |
| 5559PFC-A | 5559 | neuron | 19.8 |
| 5559PFC-B | 5559 | neuron | 19.8 |
| 5559PFC-C | 5559 | neuron | 19.8 |
| 5657-Oligo-3 | 5657 | oligodendrocyte | 82 |
| 5657-Oligo-5 | 5657 | oligodendrocyte | 82 |
| 5657-Oligo-7 | 5657 | oligodendrocyte | 82 |
| 5657PFC-A | 5657 | neuron | 82 |
| 5657PFC-B | 5657 | neuron | 82 |
| 5657PFC-C | 5657 | neuron | 82 |
| 5817-Oligo-4 | 5817 | oligodendrocyte | 0.6 |
| 5817-Oligo-6 | 5817 | oligodendrocyte | 0.6 |
| 5817-Oligo-7 | 5817 | oligodendrocyte | 0.6 |
| 5817PFC-A | 5817 | neuron | 0.6 |
| 5817PFC-B | 5817 | neuron | 0.6 |
| 5817PFC-C | 5817 | neuron | 0.6 |
| 5823-Oligo-6 | 5823 | oligodendrocyte | 82.7 |
| 5823-Oligo-7 | 5823 | oligodendrocyte | 82.7 |
| 5823-Oligo-8 | 5823 | oligodendrocyte | 82.7 |
| 5823PFC-A | 5823 | neuron | 82.7 |
| 5823PFC-C | 5823 | neuron | 82.7 |
| 5871-Oligo-4 | 5871 | oligodendrocyte | 2 |
| 5871-Oligo-5 | 5871 | oligodendrocyte | 2 |
| 5871-Oligo-7 | 5871 | oligodendrocyte | 2 |
| 5871-Neuron-5 | 5871 | neuron | 2 |
| 5871-Neuron-6 | 5871 | neuron | 2 |

| Cell ID | Depth | Technology | Tumor Grade |
|---|---|---|---|
| BOS-EA01 | 0.5X | PTA | II |
| BOS-EA02 | 0.5X | PTA | II |
| BOS-EA03 | 0.5X | PTA | II |
| BOS-EA04 | 0.5X | PTA | II |
| BOS-EA05 | 0.5X | PTA | II |
| BOS-EA06 | 0.5X | PTA | II |
| BOS-EA07 | 0.5X | PTA | II |
| BOS-EA08 | 0.5X | PTA | II |
| BOS-EA09 | 0.5X | PTA | II |
| BOS-EA10 | 0.5X | PTA | II |
| BOS-EA11 | 0.5X | PTA | II |
| BOS-EB01 | 0.5X | PTA | II |
| BOS-EB02 | 0.5X | PTA | II |
| BOS-EB03 | 0.5X | PTA | II |
| BOS-EB04 | 0.5X | PTA | II |
| BOS-EB05 | 0.5X | PTA | II |
| BOS-EB06 | 0.5X | PTA | II |
| BOS-EB07 | 0.5X | PTA | II |
| BOS-EB08 | 0.5X | PTA | II |
| BOS-EB09 | 0.5X | PTA | II |
| BOS-EB10 | 0.5X | PTA | II |
| BOS-EB11 | 0.5X | PTA | II |
| BOS-EC01 | 0.5X | PTA | II |
| BOS-EC02 | 0.5X | PTA | II |
| BOS-EC03 | 0.5X | PTA | II |
| BOS-EC04 | 0.5X | PTA | II |
| BOS-EC05 | 0.5X | PTA | II |
| BOS-EC06 | 0.5X | PTA | II |
| BOS-EC07 | 0.5X | PTA | II |
| BOS-EC08 | 0.5X | PTA | II |
| BOS-EC09 | 0.5X | PTA | II |
| BOS-EC10 | 0.5X | PTA | II |
| BOS-EC11 | 0.5X | PTA | II |
| BOS-ED01 | 0.5X | PTA | II |
| BOS-ED02 | 0.5X | PTA | II |
| BOS-ED03 | 0.5X | PTA | II |
| BOS-ED04 | 0.5X | PTA | II |
| BOS-ED05 | 0.5X | PTA | II |
| BOS-ED06 | 0.5X | PTA | II |
| BOS-ED07 | 0.5X | PTA | II |
| BOS-ED08 | 0.5X | PTA | II |
| BOS-ED09 | 0.5X | PTA | II |
| BOS-ED10 | 0.5X | PTA | II |

*Continued on next page*

Supplementary Table 3 – *Continued from previous page*

| Cell ID | Depth | Technology | Tumor Grade |
|---|---|---|---|
| BOS-ED11 | 0.5X | PTA | II |
| BOS-EE01 | 0.5X | PTA | II |
| BOS-EE02 | 0.5X | PTA | II |
| BOS-EE03 | 0.5X | PTA | II |
| BOS-EE04 | 0.5X | PTA | II |
| BOS-EE05 | 0.5X | PTA | II |
| BOS-EE06 | 0.5X | PTA | II |
| BOS-EE07 | 0.5X | PTA | II |
| BOS-EE08 | 0.5X | PTA | II |
| BOS-EE09 | 0.5X | PTA | II |
| BOS-EE10 | 0.5X | PTA | II |
| BOS-EE11 | 0.5X | PTA | II |
| BOS-EF01 | 0.5X | PTA | II |
| BOS-EF02 | 0.5X | PTA | II |
| BOS-EF03 | 0.5X | PTA | II |
| BOS-EF04 | 0.5X | PTA | II |
| BOS-EF05 | 0.5X | PTA | II |
| BOS-EF06 | 0.5X | PTA | II |
| BOS-EF07 | 0.5X | PTA | II |
| BOS-EF08 | 0.5X | PTA | II |
| BOS-EF09 | 0.5X | PTA | II |
| BOS-EF10 | 0.5X | PTA | II |
| BOS-EF11 | 0.5X | PTA | II |
| BOS-EG01 | 0.5X | PTA | II |
| BOS-EG02 | 0.5X | PTA | II |
| BOS-EG03 | 0.5X | PTA | II |
| BOS-EG04 | 0.5X | PTA | II |
| BOS-EG05 | 0.5X | PTA | II |
| BOS-EG06-BOSE-4 | 0.5X | PTA | II |
| BOS-EG06 | 0.5X | PTA | II |
| BOS-EG07 | 0.5X | PTA | II |
| BOS-EG08 | 0.5X | PTA | II |
| BOS-EG09 | 0.5X | PTA | II |
| BOS-EG10 | 0.5X | PTA | II |
| BOS-EG11 | 0.5X | PTA | II |
| BOS-EH02 | 0.5X | PTA | II |
| BOS-EH03 | 0.5X | PTA | II |
| BOS-EH04 | 0.5X | PTA | II |
| BOS-EH05 | 0.5X | PTA | II |
| BOS-EH06 | 0.5X | PTA | II |
| BOS-EH07 | 0.5X | PTA | II |
| BOS-EH08 | 0.5X | PTA | II |

*Continued on next page*

Supplementary Table 3 – *Continued from previous page*

| Cell ID | Depth | Technology | Tumor Grade |
|---|---|---|---|
| BOS-EH09 | 0.5X | PTA | II |
| BOS-EH10 | 0.5X | PTA | II |
| BOS-EH11 | 0.5X | PTA | II |
| BOS-5A01 | 0.5X | PTA | III |
| BOS-5A02 | 0.5X | PTA | III |
| BOS-5A03 | 0.5X | PTA | III |
| BOS-5A04 | 0.5X | PTA | III |
| BOS-5A05 | 0.5X | PTA | III |
| BOS-5A06 | 0.5X | PTA | III |
| BOS-5A07 | 0.5X | PTA | III |
| BOS-5A08 | 0.5X | PTA | III |
| BOS5-A09 | 0.5X | PTA | III |
| BOS-5A10 | 0.5X | PTA | III |
| BOS-5A11 | 0.5X | PTA | III |
| BOS-5A12 | 0.5X | PTA | III |
| BOS-5B01 | 0.5X | PTA | III |
| BOS-5B02 | 0.5X | PTA | III |
| BOS-5B03 | 0.5X | PTA | III |
| BOS-5B04 | 0.5X | PTA | III |
| BOS-5B05 | 0.5X | PTA | III |
| BOS-5B06 | 0.5X | PTA | III |
| BOS-5B07 | 0.5X | PTA | III |
| BOS-5B08 | 0.5X | PTA | III |
| BOS-5B09 | 0.5X | PTA | III |
| BOS-5B10 | 0.5X | PTA | III |
| BOS-5B11 | 0.5X | PTA | III |
| BOS-5B12 | 0.5X | PTA | III |
| BOS-5C01 | 0.5X | PTA | III |
| BOS-5C02 | 0.5X | PTA | III |
| BOS-5C03 | 0.5X | PTA | III |
| BOS-5C04 | 0.5X | PTA | III |
| BOS-5C05 | 0.5X | PTA | III |
| BOS-5C06 | 0.5X | PTA | III |
| BOS-5C07 | 0.5X | PTA | III |
| BOS-5C08 | 0.5X | PTA | III |
| BOS-5C09 | 0.5X | PTA | III |
| BOS-5C10 | 0.5X | PTA | III |
| BOS-5C11 | 0.5X | PTA | III |
| BOS-5C12 | 0.5X | PTA | III |
| BOS-5D01 | 0.5X | PTA | III |
| BOS-5D02 | 0.5X | PTA | III |
| BOS-5D03 | 0.5X | PTA | III |

*Continued on next page*

Supplementary Table 3 – *Continued from previous page*

| Cell ID | Depth | Technology | Tumor Grade |
|---|---|---|---|
| BOS-5D04 | 0.5X | PTA | III |
| BOS-5D05 | 0.5X | PTA | III |
| BOS-5D06 | 0.5X | PTA | III |
| BOS-5D07 | 0.5X | PTA | III |
| BOS-5D09 | 0.5X | PTA | III |
| BOS-5D10 | 0.5X | PTA | III |
| BOS-5D11 | 0.5X | PTA | III |
| BOS-5E01 | 0.5X | PTA | III |
| BOS-5E02 | 0.5X | PTA | III |
| BOS-5E03 | 0.5X | PTA | III |
| BOS-5E04 | 0.5X | PTA | III |
| BOS-5E05 | 0.5X | PTA | III |
| BOS-5E06 | 0.5X | PTA | III |
| BOS-5E07 | 0.5X | PTA | III |
| BOS-5E08 | 0.5X | PTA | III |
| BOS-5E09 | 0.5X | PTA | III |
| BOS-5E10 | 0.5X | PTA | III |
| BOS-5E11 | 0.5X | PTA | III |
| BOS-5F01 | 0.5X | PTA | III |
| BOS-5F02 | 0.5X | PTA | III |
| BOS-5F03 | 0.5X | PTA | III |
| BOS-5F04 | 0.5X | PTA | III |
| BOS-5F05 | 0.5X | PTA | III |
| BOS-5F06 | 0.5X | PTA | III |
| BOS-5F07 | 0.5X | PTA | III |
| BOS-5F08 | 0.5X | PTA | III |
| BOS-5F09 | 0.5X | PTA | III |
| BOS-5F10 | 0.5X | PTA | III |
| BOS-5F11 | 0.5X | PTA | III |
| BOS-5G01 | 0.5X | PTA | III |
| BOS-5G02 | 0.5X | PTA | III |
| BOS-5G03 | 0.5X | PTA | III |
| BOS-5G04 | 0.5X | PTA | III |
| BOS-5G05 | 0.5X | PTA | III |
| BOS-5G06 | 0.5X | PTA | III |
| BOS-5G07 | 0.5X | PTA | III |
| BOS-5G08 | 0.5X | PTA | III |
| BOS-5G09 | 0.5X | PTA | III |
| BOS-5G10-BOS5-3 | 0.5X | PTA | III |
| BOS-5G10 | 0.5X | PTA | III |
| BOS-5G11 | 0.5X | PTA | III |
| BOS-5H01 | 0.5X | PTA | III |

*Continued on next page*

Supplementary Table 3 – *Continued from previous page*

| Cell ID | Depth | Technology | Tumor Grade |
|---|---|---|---|
| BOS-5H02 | 0.5X | PTA | III |
| BOS-5H03 | 0.5X | PTA | III |
| BOS-5H04 | 0.5X | PTA | III |
| BOS-5H05 | 0.5X | PTA | III |
| BOS-5H06 | 0.5X | PTA | III |
| BOS-5H07 | 0.5X | PTA | III |
| BOS-5H08 | 0.5X | PTA | III |
| BOS-5H09 | 0.5X | PTA | III |
| BOS-5H10 | 0.5X | PTA | III |
| BOS-5H11 | 0.5X | PTA | III |
| BOS-E-BrainBulk | 55X | bulk | II |
| BOS5-BrainBulk | 55X | bulk | III |
| BOS-EBlood | 25X | bulk | II |

**Supplementary Table 3**: Samples analyzed in the meningioma scWGS analysis. All of the single cells were amplified with PTA. "BOS-E" and "BOS-5" prefixes are equivalent to grade II and III, respectively. Tumors of both grades came from the same donor.
